# Leveraging cell type-specificity for gene set analysis of single cell transcriptomics

**DOI:** 10.1101/2024.09.25.615040

**Authors:** H. Robert Frost

## Abstract

Although single cell RNA-sequencing (scRNA-seq) provides unprecedented insights into the biology of complex tissues, analyzing such data on a gene-by-gene basis is challenging due to the large number of tested hypotheses and consequent low statistical power and difficult interpretation. These issues are magnified by the increased noise, significant sparsity and multi-modal distributions characteristic of single cell data. One promising approach for addressing these challenges is gene set testing, or pathway analysis. Unfortunately, statistical and biological differences between single cell and bulk transcriptomic data make it challenging to use existing gene set collections, which were developed for bulk tissue analysis, on scRNA-seq data. In this paper, we describe a procedure for customizing gene set collections originally created for bulk tissue analysis to reflect the structure of gene activity within specific cell types. Our approach leverages information about mean gene expression in the 81 human cell types profiled via scRNA-seq by the Human Protein Atlas (HPA) Single Cell Type Atlas. This HPA information is used to compute cell type-specific gene and gene set weights that can be used to filter or weight gene set collections. As demonstrated through the analysis of immune cell scRNA-seq data using gene sets from the Molecular Signatures Database (MSigDB), accounting for cell type-specificity can significantly improve gene set testing power and interpretability. An example vignette along with gene and gene set weights for the 81 HPA SCTA cell types and the MSigDB collections are available at https://hrfrost.host.dartmouth.edu/SCGeneSetOpt/.

## 1 Introduction

### 1.1 Single cell analysis challenges

Although single cell assays such as single cell RNA-sequencing (scRNA-seq) [1] are a powerful tool for studying complex tissues, technical and biological limitations make statistical analysis challenging [2, 3]. Single cell methods profile very small amounts of genomic material, leading to significant amplification bias and sparsity relative to bulk tissue assays [4]. Single cell approaches for quality control, normalization and statistical analysis (e.g., zero-inflated models) only partially address these challenges [5, 6]. In addition to the challenges of noise and sparsity, important biological differences exist between bulk tissue and single cell data. As the average over a large number of cells, bulk tissue measurements are non-sparse, typically unimodal and, in many cases, approximately normally distributed. In contrast, single cell datasets reflect a heterogenous mixture of cell types and states resulting in multi-modal and non-normal distributions [4]. The diverse mixture of cell types and states found in complex tissues also leads to significant differences in gene expression patterns between bulk tissue and single cell data. As evidenced by projects such as the Human Protein Atlas (HPA) [7], gene activity in bulk tissue, as quantified via gene expression or protein abundance, can differ substantially from the activity occurring within the cell subpopulations comprising the tissue.

The HPA repository was recently updated with the Single Cell Type Atlas (SCTA) [8], which captures gene expression values for 81 common human cell types as measured by scRNA-seq on healthy tissue for 31 different tissue types. For the HPA SCTA, the source scRNA-seq data was obtained from the Single Cell Expression Atlas (SCEA) [9], the Human Cell Atlas (HCA) [10, 11], the Gene Expression Omnibus (GEO) [12], and the European Genome-phenome Archive (EGA) [13]. The datasets from these source repositories were carefully curated to identify high-quality scRNA-seq data measured on 31 different tissue types where the samples were obtained from healthy individuals and processed without cell type enrichment. Using this data, cell type clusters where identified representing 81 distinct cell types. The mean expression profile of each cell type enables the quantification of the cell type-specificity of human protein coding genes.

Importantly, mean gene expression differs not only between different cell types but also between bulk tissue samples and the cell types that comprise that tissue. Figure 1 illustrates these marginal differences for a subset of genes in the Molecular Signatures Database (MSigDB) [14] Hallmark TGF-*β* signaling pathway based on the cell types captured via scRNA-seq in human skin [15] as represented in the HPA SCTA. As illustrated by this figure, gene expression values measured via scRNA-seq on the cell types that comprise skin can look very different from the values computed via bulk RNA-seq on skin samples, which will be a weighted average of the cell type-specific measurements. The pattern of coexpression can also vary significantly between single cells and bulk tissue. A comparison of gene co-expression in single cell and bulk glioblastoma samples performed by Wang et al. [16] found that over 90% of the gene co-expression pairs were unique to either the bulk or single cell data. This dramatic difference in the pattern of coexpression is due to the fact that co-expression at the bulk tissue level is often driven by variation in the proportion of cell types in a given tissue which can bear little resemblance to gene co-expression across cells of a specific type [16, 17]. Genes that are uncorrelated at the single cell level can appear to be correlated at the bulk tissue level if the mean expression varies across cell types and cell type proportions vary across bulk samples. The inverse is also possible, i.e., genes whose expression is correlated at the single cell level can appear uncorrelated in bulk tissue.

**Figure 1:**
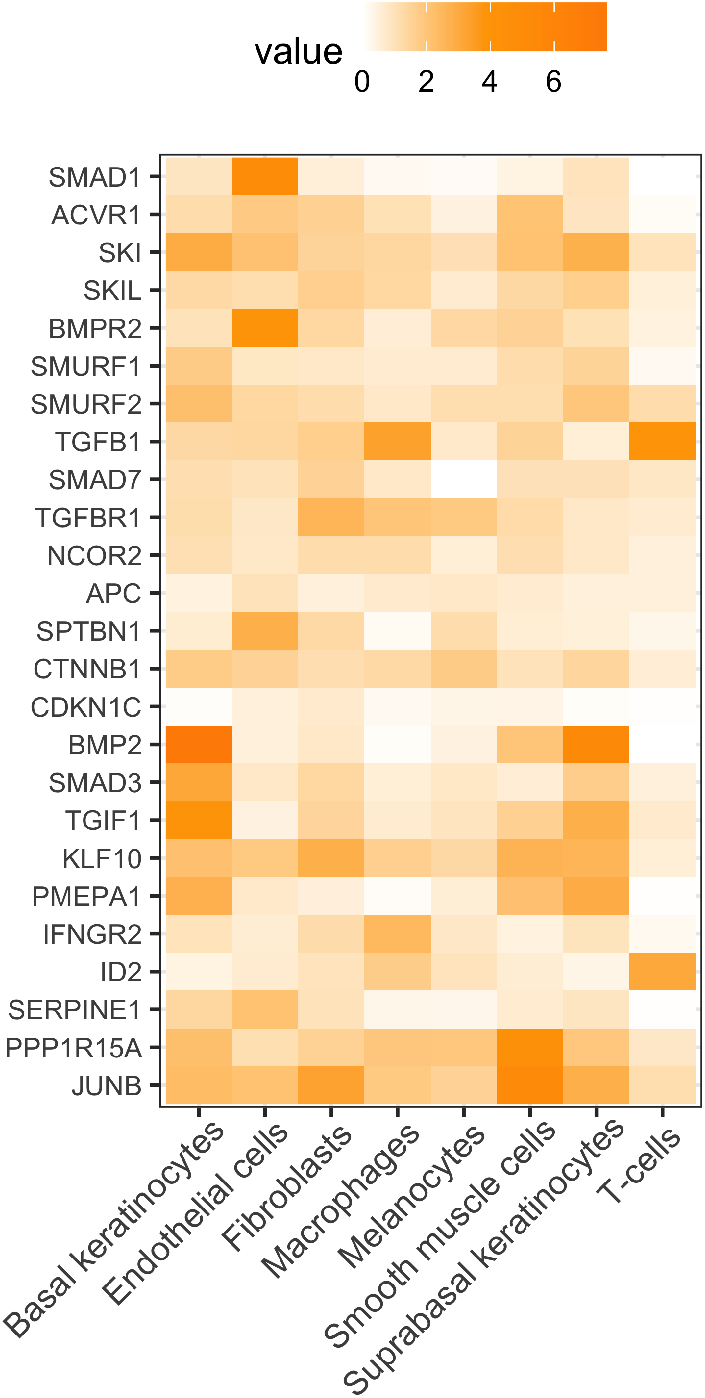
Cell type-specific expression of genes in the MSigDB Hallmark TGF-*β* signaling pathway. Cells represent the fold-change in mean expression of the gene in a given cell type relative to the average across all 81 cell types profiled by the HPA Single Cell Type Atlas.

A similar issue exists for the association of gene expression values with a given experimental condition, i.e., the associations found at the bulk tissue level can be very dissimilar to those found for a specific cell type. Figure 2 provides a simplified illustration of the marginal and joint distribution characteristics of single cell and bulk tissue expression data. In this figure, the marginal distribution is represented by density plots for a single gene while the joint distribution is represented by covariance matrices. Collectively, the distributional differences between single cell and bulk tissue genomic data make it challenging to successfully analyze single cell expression data using biological models originally developed for bulk tissue, which represent the pattern of gene product abundance within an average cell.

**Figure 2:**
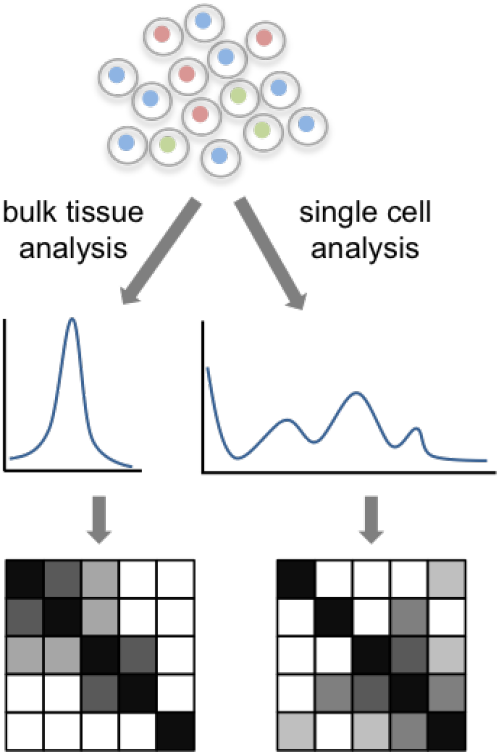
Bulk tissue vs. single cell distributions. The middle and bottom rows illustrate approximate marginal and joint expression distributions.

### 1.2 Gene set testing for single cell data

Although high-dimensional genomic data provides a molecular-level lens on biological systems, the gain in fidelity obtained by testing thousands of genomic variables comes at the price of impaired interpretation, loss of power due to multiple hypothesis correction and poor reproducibility [18, 19]. To help address these challenges for bulk tissue data, researchers developed gene set testing, or pathway analysis, methods [20]. Gene set testing is an effective hypothesis aggregation technique that lets researchers step back from the level of individual genomic variables and explore associations for biologically meaningful groups of genes. Focusing on a small number of pathways can substantially improve power, interpretation and replication relative to an analysis focused on individual genomic variables [21]. The benefits that gene set-based hypothesis aggregation offers for the analysis of bulk tissue data are even more pronounced for single cell data given increased technical variance and sparsity. Although significant progress has been made developing gene set testing methods, including methods developed by us that are specifically optimized for scRNA-seq data [22,23], and building and maintaining gene set collections, existing collections were largely developed for the analysis of bulk tissue data. This is problematic since many gene sets in collections like the Molecular Signatures Database (MSigDB) [14] are defined to contain groups of genes whose expression in bulk tissue is correlated (e.g., MSigDB cancer modules [24]) or is associated with a specific experimental variable (e.g., MSigDB chemical and genetic perturbations). Such gene sets will often represent biological associations that do not hold at the single cell level [16]. It is important to note that new collections, e.g., the Human Cell Atlas-based MSigDB C8 collection [25], are being developed that contain gene sets derived from scRNA-seq data.

## 2 Data and methods

To address the bulk tissue bias that exists in most public gene set collections, we have developed a procedure for customizing gene set collections to reflect the structure of gene activity within specific cell types as measured by single cell transcriptomic assays. Our approach leverages information about mean gene expression in the 81 human cell types profiled via scRNA-seq by the HPA SCTA. As detailed below, this SCTA information was used to compute cell type-specific gene and gene set weights that can be used to filter or weight gene set collections. An example vignette and gene and gene set weights for the 81 HPA SCTA cell types and MSigDB collections are available at https://hrfrost.host.dartmouth.edu/SCGeneSetOpt/.

### 2.1 Data sources

The following data sources were leveraged to compute the cell type-specific gene and gene set weights and generate the paper results:

- Human Protein Atlas Single Cell Type Atlas (HPA SCTA): Information about the cell type-specific expression of human protein coding genes was obtained from the HPA SCTA via the downloadable file https://www.proteinatlas.org/download/rna_single_cell_type.tsv.zip.
- Molecular Signatures Database (MSigDB): Gene set definitions were obtained from the MSigDB v2024.1 via the downloadable files at https://www.gsea-msigdb.org/gsea/msigdb/index.jsp.
- 10k PBMC3k scRNA-seq data: The PBMC scRNA-seq dataset used to generated the results in Section 3 is also used in the Seurat Guided Clustering Tutorial (https://satijalab.org/seurat/articles/pbmc3k_tutorial.html), is included in the *SeuratData* R package and is freely accessible from 10x Genomics via a Creative Commons Attribution license at https://cf.10xgenomics.com/samples/cell/pbmc3k/pbmc3k_filtered_gene_bc_matrices.tar.gz. Processing of the PBMC3k dataset was performed using the same logic employed in the Seurat Guided Clustering Tutorial, which is also contained in the vignette available at https://hrfrost.host.dartmouth.edu/SCGeneSetOpt/.

### 2.2 Computation of cell type-specific gene and gene set weights

Building on our prior work creating customized versions of MSigDB for different normal human tissue types [26], our method first computes cell type-specific weights for each protein coding human gene for all 81 normal human cell types profiled by the HPA SCTA (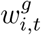 for gene *i* in cell type *t*). Specifically, 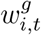 is set to the fold-change between the mean normalized transcript abundance of gene *i* in cell type *t* as computed via scRNA-seq relative to the average across all 81 cell types These gene-level weights are then leveraged to compute cell type-specific gene set weights for the sets in the MSigDB collections. Specifically, a weight 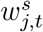 for each gene set *j* in a target collection is computed as the -log of the p-value from a one-sided, two-sample test t test comparing the 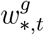 values for the genes in set *j* to the 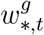 for genes not in *j* (this is similar to the competitive gene set test implemented by the *cameraPR* method in the R limma package [27]).

### 2.3 Using gene weights for annotation filtering and weighting

The cell type-specific gene weights 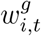 can be used to customize gene sets via either annotation filtering or weighting. Gene set annotations can be customized for cell type *t* by simply removing all annotations for each gene *i* if 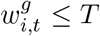, where *T* is a threshold (*T ≥* 0) that can be user specified or selected to optimize a gene set testing performance metric, e.g., replication of gene set testing results across related datasets. Filtering has the benefit of parsimony (it results in smaller and more easily intepreted sets), improved power (the remaining set members should be more likely to capture the relevant biological signal), and can work with any gene set testing method, however, it does require the specification of a threshold and ignores most of the information contained in the continuous weights. An alternate approach that uses the continuous gene weights and does not require a threshold is annotation weighting. In this scenario, the log of the gene weights is used to provide a directional weight that can be used with gene set testing methods like *fry* (see Section 2.6 for more details) that accept gene weights.

### 2.4 Using gene set weights for collection filtering and weighting

Similar to the application of gene-level weights, the cell type-specific gene set weights 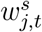 can be used for either filtering or weighting. Filtering can be performed by removing (or not using) all sets in a given collection where 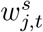 is less than some threshold. Collection filtering has the benefits of interpretation and statistical power since a smaller number of more biologically relevant gene sets are tested, which makes interpretation easier and reduces the multiple hypothesis correction penalty. Like annotation filtering, the downsides of collection filtering include the need for a specific threshold and fact that most of the information in the weights is not used. The weights can alternatively be used for p-value weighting (e.g., weighted FDR [28]) following gene set testing, which avoids the need for a specific threshold.

### 2.5 Choice of cell type weights

The effectiveness of the filtering and weighting techniques detailed in Sections 2.3 and 2.4 is strongly dependent on the what cell type weight is employed. Considerations for several common scenarios are discussed below:

- *Analysis of a single cell type in different experimental conditions* : For this scenario, gene set analysis is typically performed to identify sets that are differentially active between the experimental conditions. Using gene and/or gene set weights for type of cell in the dataset is usually appropriate since this will prioritize the pathways/functions most critical to the biology of that cell type.
- *Analysis of multiple cell types in different experimental conditions* : For this scenario, the goal of gene set analysis is usually to identify sets that are differentially active between the experimental conditions irrespective of the cell type. In this case, using an average (or weighted average) of the weights for all cell types present in the data can be effective. Similar to the single cell type case, this will priorize the pathways/functions most relevant to the function of those cell types.
- *Comparative analysis of different cell types in the same experimental condition* : For this scenario, gene set analysis aims to identify sets that are differentially active in cells of one type relative to cells of a different type. If the analysis is primarily focused on one of the cell types, then the weights for that type could be used. Alternatively, the average of the weights (or maximum weight) for the analyzed cell types could be used.

The results in Section 3 correspond to the last scenario and weights for just one of the cell types in the comparison are used (either B cell or T cell weights).

### 2.6 Gene set testing

The gene set testing results shown in Section 3 were generated using two techniques, both availble via the *limma* R package [27], for population-level gene set analysis:

- *Camera* [27]: This technique implements a competitive and population-level gene set test that accounts for inter-gene correlation.
- *Fry* [29]: This technique implementes a self-contained and population-level gene set test that can accept gene-level weights.

Both methods were executed using default parameters unless otherwise specified.

## 3 Results

To evaluate the cell type-specific weight model detailed in Section 2.2, we computed gene and gene set weights for all 81 normal human cell types included in the HPA SCTA. For each of these cell types, gene weights were generated for all human protein coding genes profiled by the HPA SCTA and gene set weights were calculated for all collections in v2024.1 of the MSigDB. To illustrate the utility of these weights and their application for the filtering and weighting approaches detailed in Sections 2.3 and 2.4, we performed various gene set analyses of the 10x PBMC3k example scRNA-seq dataset (visualized in Figure 3) using both B and T cell weights and the sets in the Gene Ontology Biological Process collection (MSigDB collection C5.GO.BP). Results from these analyses are detailed in Sections 3.1-3.4 below.

**Figure 3:**
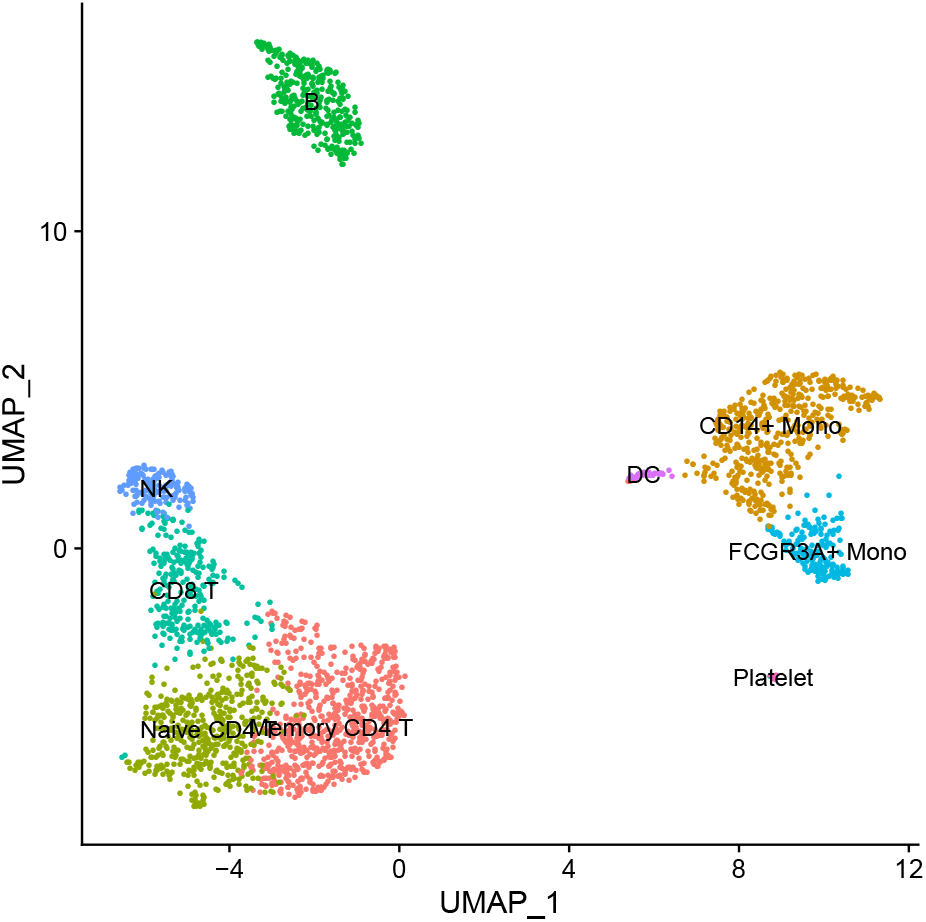
Projection of PBMC scRNA-seq data onto the first two UMAP dimensions. Each point in the plot represents one cell.

Both the cell type-specific weights and R logic for the B cell-based results are available on the paper website (https://hrfrost.host.dartmouth.edu/SCGeneSetOpt/).

### 3.1 Collection filtering using B cell gene set weights

As illustrated by Table 1, which lists the top ten MSigDB C5.GO.BP (Gene Ontology Biological Process) terms for B cells, the gene sets with the largest *w*^*s*^ values clearly reflect the known biological features of B cells.

**Table 1:** Top 10 MSigDB C5.GO.BP gene sets based on B cell gene set weights.

| Gene set | Weight ($w^s$) |
| --- | --- |
| GOBP'B'CELL'RECEPTOR'SIGNALING'PATHWAY | 412 |
| GOBP'B'CELL'ACTIVATION | 270 |
| GOBP'ANTIGEN'RECEPTOR'MEDIATED'SIGNALING'PATHWAY | 206 |
| GOBP'ADAPTIVE'IMMUNE'RESPONSE | 205 |
| GOBP'LYMPHOCYTE'ACTIVATION | 200 |
| GOBP'B'CELL'PROLIFERATION | 186 |
| GOBP'IMMUNE'RESPONSE'REGULATING'CELL'SURFACE'RECEPTOR'SIGNALING'PATHWAY | 185 |
| GOBP'CELL'ACTIVATION | 161 |
| GOBP'POSITIVE'REGULATION'OF'IMMUNE'RESPONSE | 155 |
| GOBP'IMMUNE'RESPONSE'REGULATING'SIGNALING'PATHWAY | 155 |

Following the approach outlined in Section 2.4, we used the B cell-based gene set weights to filter the MSigDB C5.GO.BP collection with the goal of both improving statistical power by reducing the multiple hypothesis correction burden and improving the biological relevance and interpretability of the analysis by only testing sets specific to B cells. To perform this analysis, we created a filtered version of the C5.GO.BP collection that retained the sets with B cell weights above 10, which corresponds to 4.5% of the 5,777 C5.GO.BP sets retained after alignment with the PBMC scRNA-seq genes (annotations were removed for genes not captured in the PBMC data and then sets were eliminated if they had less then 5 or greater than 200 members). We then performed a population-level and competitive gene set analysis using the *camera* method comparing set expression among B cells against expression in non-B cells. Weight-based collection filtering had the desired effect of improving the multiple correction-adjusted statistical significance of the results. Specifically, without filtering only 36 terms had FDR values *<* 0.1; with filtering this number increased to 67. This trend is visualized in Figure 4.

**Figure 4:**
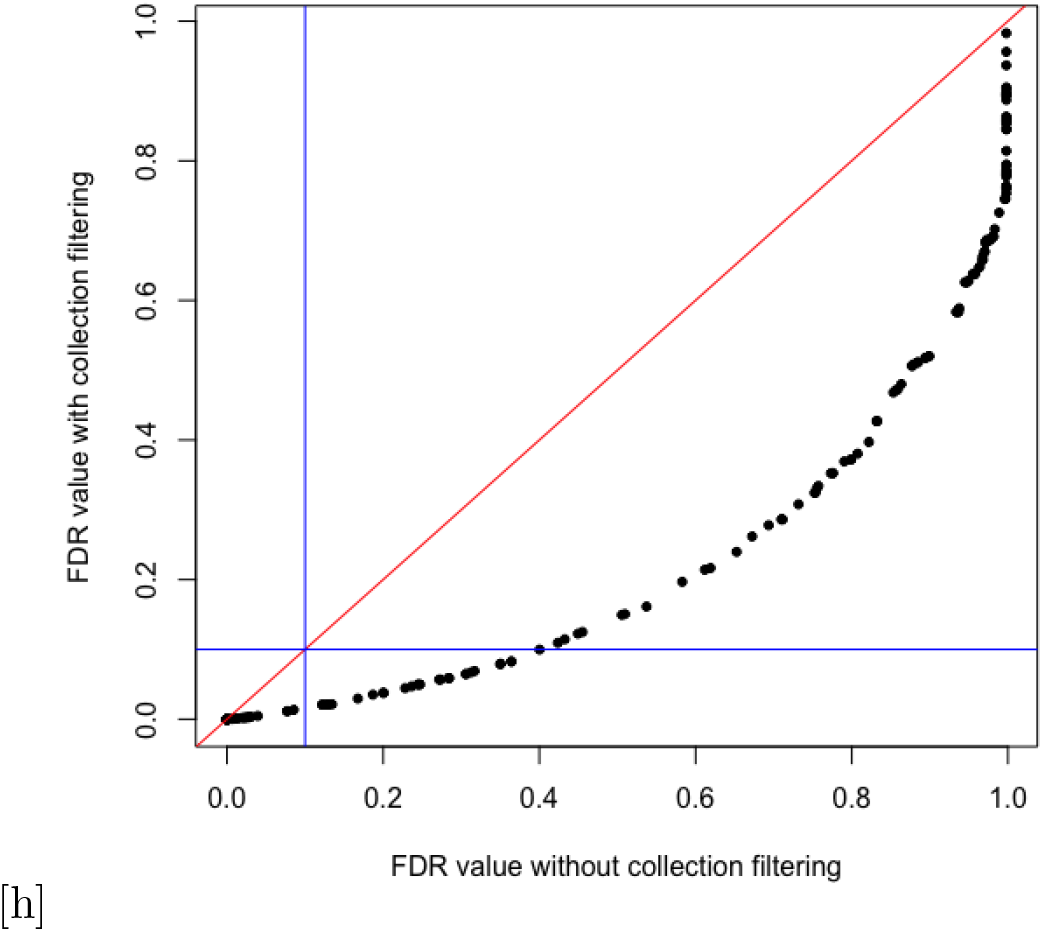
Distribution of FDR values from a gene set analysis comparing B cells against other cell types in the PBMC data using the MSigDB C5.GO.BP collection. Each point captures the FDR values for one of the terms remaining after collection filtering with the x-axis representing the non-filtered FDR value and the y-axis representing the filtered FDR value. Blue lines reflect the FDR threshold of 0.1 and the red line reflects expected distribution for equal FDR values.

### 3.2 Collection filtering using T cell gene set weights

Table 2 lists the top ten MSigDB C5.GO.BP (Gene Ontology Biological Process) terms according to the T cell-specific gene set weights. Similar to the top terms for B cells shown in Table 1, the Gene Ontology terms with the largest *w*^*s*^ values effectively capture the key aspects of T cell biology.

**Table 2:** Top 10 MSigDB C5.GO.BP gene sets based on T cell gene set weights.

| Gene set | Weight ($w^s$) |
| --- | --- |
| GOBP'ADAPTIVE'IMMUNE'RESPONSE | 500.0 |
| GOBP'T'CELL'ACTIVATION | 109 |
| GOBP'T'CELL'RECEPTOR'SIGNALING'PATHWAY | 101 |
| GOBP'LYMPHOCYTE'ACTIVATION | 91 |
| GOBP'ANTIGEN'RECEPTOR'MEDIATED'SIGNALING'PATHWAY | 81 |
| GOBP'BIOLOGICAL'PROCESS'INVOLVED'IN'INTERSPECIES'INTERACTION'BETWEEN'ORGANISMS | 75 |
| GOBP'T'CELL'DIFFERENTIATION | 69 |
| GOBP'ALPHA'BETA'T'CELL'ACTIVATION | 68 |
| GOBP'POSITIVE'REGULATION'OF'IMMUNE'SYSTEM'PROCESS | 68 |
| GOBP'CELL'ACTIVATION | 67 |

Similar to the B cell analysis above, we created a filtered version of the C5.GO.BP collection that retained the sets with T cell weights above 10, which corresponds to 4.4% of the 5,777 C5.GO.BP sets retained after alignment with the PBMC scRNAseq genes. We then performed a gene set analysis using the *camera* method comparing set expression among T cells against expression in nonT cells. Weight-based collection filtering again had the desired effect of improving the multiple correction-adjusted statistical significance of the results. Specifically, without filtering 56 terms had FDR values *<* 0.1, with filtering this number increased to 68. This trend is visualized in Figure 5.

**Figure 5:**
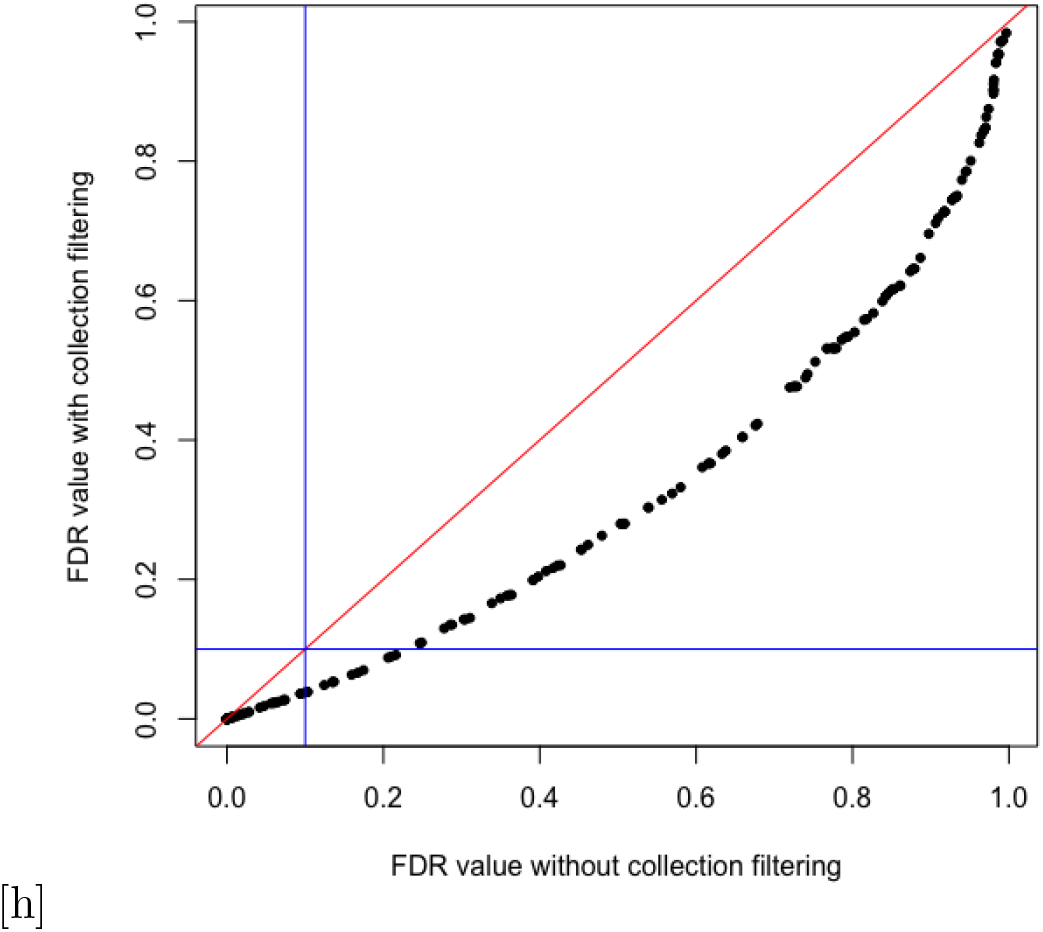
Distribution of FDR values from a gene set analysis comparing T cells against other cell types in the PBMC data using the MSigDB C5.GO.BP collection. Each point captures the FDR values for one of the terms remaining after collection filtering with the x-axis representing the non-filtered FDR value and the y-axis representing the filtered FDR value. Blue lines reflect the FDR threshold of 0.1 and the red line reflects expected distribution for equal FDR values.

### 3.3 Annotation weighting using B cell gene weights

Following the approach in Section 2.3, we used the B cell-based gene weights to perform a weighted gene set analysis using the *fry* method. Specifically, set expression in B cells was compared to set expression in non-B cells and this analysis was performed both without weights and with gene weights set to the log2 of the B cell-based gene weight plus a pseudo-count of 1e-4 (the log transformation generates sign-based directional weights as need by *fry*). This weighting scheme prioritizes genes that are strongly up-regulated or down-regulated in B cells according to the HPA SCTA. The *fry* method (which is a fast approximation of the *roast* technique) generated unexpectedly small FDR values (much smaller than *camera*), however, the rank ordering of the sets based on significance was similar to *camera* and matched the expected biology of B cells. Use of B cell gene weights improved the statistical power of the gene set analysis. Specifically, without weights 1,965 terms had FDR values *<* 1*e −* 4, with weights this number increased to 2,348.

### 3.4 Annotation weighting using T cell gene weights

Similar to the B cell analysis in Section 3.3, we used T cell-based gene weights to perform a weighted gene set analysis comparing set expression in T cells to expression in non-T cells. Use of T cell gene weights also improved the statistical power of the gene set analysis. Specifically, without weights 2,920 terms had FDR values *<* 1*e −* 4, with weights this number increased to 3,590.

## 4 Conclusions

Gene set testing is a powerful tool for the analysis of scRNA-seq data that addresses the challenges of sparsity and noise. Unfortunately, the utility of gene set testing for single cell data is limited by the fact that most existing gene set collections were developed to capture gene activity within bulk tissue data, which can differ substantially from gene activity in specific cell types. In particular, the pattern of gene co-expression found among cells of a specific type is often significantly different from the pattern seen in bulk tissue samples, which is driven by cell type proportions. A similar issue exists for the association of gene expression values with a given experimental condition. To address this challenge, we explored methods for computing gene and gene set weights using information about the cell type-specificity of human protein coding genes from the HPA SCTA. These cell type-specific weights can be leveraged to improve the power and interpretability of gene set analyses through the filtering or weighting of gene set collections or gene set annotations. To support this type of analysis by other researchers, an example vignette along with gene and gene set weights for the 81 HPA SCTA cell types and MSigDB collections are available at https://hrfrost.host.dartmouth.edu/SCGeneSetOpt/.

## References

[1] Evan Z Macosko, Anindita Basu, Rahul Satija, James Nemesh, Karthik Shekhar, Melissa Goldman, Itay Tirosh, Allison R Bialas, Nolan Kamitaki, Emily M Martersteck, John J Trombetta, David A Weitz, Joshua R Sanes, Alex K Shalek, Aviv Regev, and Steven A McCarroll. Highly parallel genome-wide expression profiling of individual cells using nanoliter droplets. Cell, 161(5):1202–1214, May 2015.

[2] Guo-Cheng Yuan, Long Cai, Michael Elowitz, Tariq Enver, Guoping Fan, Guoji Guo, Rafael Irizarry, Peter Kharchenko, Junhyong Kim, Stuart Orkin, John Quackenbush, Assieh Saadatpour, Timm Schroeder, Ramesh Shivdasani, and Itay Tirosh. Challenges and emerging directions in single-cell analysis. Genome Biol, 18(1):84, May 2017.

[3] Lukas Heumos, Anna C Schaar, Christopher Lance, Anastasia Litinetskaya, Felix Drost, Luke Zappia, Malte D Lücken, Daniel C Strobl, Juan Henao, Fabiola Curion, Single-cell Best Practices Consortium, Herbert B Schiller, and Fabian J Theis. Best practices for single-cell analysis across modalities. Nat Rev Genet, 24(8):550–572, Aug 2023.

[4] Rhonda Bacher and Christina Kendziorski. Design and computational analysis of single-cell rna-sequencing experiments. Genome Biol, 17:63, Apr 2016.

[5] Davis J McCarthy, Kieran R Campbell, Aaron T L Lun, and Quin F Wills. Scater: pre-processing, quality control, normalization and visualization of single-cell rna-seq data in r. Bioinformatics, 33(8):1179–1186, Apr 2017.

[6] Xiaojie Qiu, Andrew Hill, Jonathan Packer, Dejun Lin, Yi-An Ma, and Cole Trapnell. Single-cell mrna quantification and differential analysis with census. Nat Methods, 14(3):309–315, Mar 2017.

[7] Mathias Uhlén, Linn Fagerberg, Björn M Hallström, Cecilia Lindskog, Per Oksvold, Adil Mardinoglu, Åsa Sivertsson, Caroline Kampf, Evelina Sjöstedt, Anna Asplund, IngMarie Olsson, Karolina Edlund, Emma Lundberg, Sanjay Navani, Cristina Al-Khalili Szigyarto, Jacob Odeberg, Dijana Djureinovic, Jenny Ottosson Takanen, Sophia Hober, Tove Alm, Per-Henrik Edqvist, Holger Berling, Hanna Tegel, Jan Mulder, Johan Rockberg, Peter Nilsson, Jochen M Schwenk, Marica Hamsten, Kalle von Feilitzen, Mattias Forsberg, Lukas Persson, Fredric Johansson, Martin Zwahlen, Gunnar von Heijne, Jens Nielsen, and Fredrik Pontén. Proteomics. tissue-based map of the human proteome. Science, 347(6220):1260419, Jan 2015.

[8] Peter J Thul, Lovisa Åkesson, Mikaela Wiking, Diana Mahdessian, Aikaterini Geladaki, Hammou Ait Blal, Tove Alm, Anna Asplund, Lars Björk, Lisa M Breckels, Anna Bäckström, Frida Danielsson, Linn Fagerberg, Jenny Fall, Laurent Gatto, Christian Gnann, Sophia Hober, Martin Hjelmare, Fredric Johansson, Sunjae Lee, Cecilia Lindskog, Jan Mulder, Claire M Mulvey, Peter Nilsson, Per Oksvold, Johan Rockberg, Rutger Schutten, Jochen M Schwenk, Åsa Sivertsson, Evelina Sjöstedt, Marie Skogs, Charlotte Stadler, Devin P Sullivan, Hanna Tegel, Casper Winsnes, Cheng Zhang, Martin Zwahlen, Adil Mardinoglu, Fredrik Pontén, Kalle von Feilitzen, Kathryn S Lilley, Mathias Uhlén, and Emma Lundberg. A subcellular map of the human proteome. Science, 356(6340), 05 2017.

[9] Irene Papatheodorou, Pablo Moreno, Jonathan Manning, Alfonso Muñoz-Pomer Fuentes, Nancy George, Silvie Fexova, Nuno A Fonseca, Anja Füllgrabe, Matthew Green, Ni Huang, Laura Huerta, Haider Iqbal, Monica Jianu, Suhaib Mohammed, Lingyun Zhao, Andrew F Jarnuczak, Simon Jupp, John Marioni, Kerstin Meyer, Robert Petryszak, Cesar Augusto Prada Medina, Carlos Talavera-López, Sarah Teichmann, Juan Antonio Vizcaino, and Alvis Brazma. Expression atlas update: from tissues to single cells. Nucleic Acids Res, 48(D1):D77–D83, 01 2020.

[10] Orit Rozenblatt-Rosen, Michael J T Stubbington, Aviv Regev, and Sarah A Teichmann. The human cell atlas: from vision to reality. Nature, 550(7677):451–453, Oct 2017.

[11] Chung-Chau Hon, Jay W Shin, Piero Carninci, and Michael J T Stubbington. The human cell atlas: Technical approaches and challenges. Brief Funct Genomics, Oct 2017.

[12] Tanya Barrett, Stephen E Wilhite, Pierre Ledoux, Carlos Evangelista, Irene F Kim, Maxim Tomashevsky, Kimberly A Marshall, Katherine H Phillippy, Patti M Sherman, Michelle Holko, Andrey Yefanov, Hyeseung Lee, Naigong Zhang, Cynthia L Robertson, Nadezhda Serova, Sean Davis, and Alexandra Soboleva. Ncbi geo: archive for functional genomics data sets–update. Nucleic Acids Res, 41(Database issue):D991–5, Jan 2013.

[13] Ilkka Lappalainen, Jeff Almeida-King, Vasudev Kumanduri, Alexander Senf, John Dylan Spalding, Saif Ur-Rehman, Gary Saunders, Jag Kandasamy, Mario Caccamo, Rasko Leinonen, Brendan Vaughan, Thomas Laurent, Francis Rowland, Pablo Marin-Garcia, Jonathan Barker, Petteri Jokinen, Angel Carreño Torres, Jordi Rambla de Argila, Oscar Martinez Llobet, Ignacio Medina, Marc Sitges Puy, Mario Alberich, Sabela de la Torre, Arcadi Navarro, Justin Paschall, and Paul Flicek. The european genome-phenome archive of human data consented for biomedical research. Nat Genet, 47(7):692–5, Jul 2015.

[14] Arthur Liberzon, Aravind Subramanian, Reid Pinchback, Helga Thorvaldsdóttir, Pablo Tamayo, and Jill P Mesirov. Molecular signatures database (msigdb) 3.0. Bioinformatics, 27(12):1739–40, Jun 2011.

[15] Llorenç Solé-Boldo, Günter Raddatz, Sabrina Schütz, Jan-Philipp Mallm, Karsten Rippe, Anke S Lonsdorf, Manuel Rodríguez-Paredes, and Frank Lyko. Single-cell transcriptomes of the human skin reveal age-related loss of fibroblast priming. Commun Biol, 3(1):188, 04 2020.

[16] Jie Wang, Shuli Xia, Brian Arand, Heng Zhu, Raghu Machiraju, Kun Huang, Hongkai Ji, and Jiang Qian. Single-cell co-expression analysis reveals distinct functional modules, co-regulation mechanisms and clinical outcomes. PLoS Comput Biol, 12(4):e1004892, Apr 2016.

[17] Megan Crow, Anirban Paul, Sara Ballouz, Z Josh Huang, and Jesse Gillis. Exploiting single-cell expression to characterize co-expression replicability. Genome Biol, 17:101, May 2016.

[18] David B Allison, Xiangqin Cui, Grier P Page, and Mahyar Sabripour. Microarray data analysis: from disarray to consolidation and consensus. Nature Reviews Genetics, 7(1):55–65, Jan 2006.

[19] Jelle J. Goeman and Peter Buehlmann. Analyzing gene expression data in terms of gene sets: methodological issues. Bioinformatics, 23(8):980–987, APR 15 2007.

[20] Purvesh Khatri, Marina Sirota, and Atul J Butte. Ten years of pathway analysis: current approaches and outstanding challenges. PLoS Computational Biology, 8(2):e1002375, Feb 2012.

[21] Aravind Subramanian, Pablo Tamayo, Vamsi K. Mootha, Sayan Mukherjee, Benjamin L. Ebert, Michael A. Gillette, Amanda Paulovich, Scott L. Pomeroy, Todd R. Golub, Eric S. Lander, and Jill P. Mesirov. Gene set enrichment analysis: A knowledge-based approach for interpreting genome-wide expression profiles. Proc Natl Acad Sci U S A, 102(43):15545–15550, October 2005.

[22] Hildreth Robert Frost. Variance-adjusted mahalanobis (vam): a fast and accurate method for cell-specific gene set scoring. Nucleic Acids Res, Jul 2020.

[23] H Robert Frost. Reconstruction set test (reset): A computationally efficient method for single sample gene set testing based on randomized reduced rank reconstruction error. PLoS Comput Biol, 20(4):e1012084, Apr 2024.

[24] Eran Segal, Nir Friedman, Daphne Koller, and Aviv Regev. A module map showing conditional activity of expression modules in cancer. Nat Genet, 36(10):1090–8, Oct 2004.

[25] Aviv Regev, Sarah A Teichmann, Eric S Lander, Ido Amit, Christophe Benoist, Ewan Birney, Bernd Bodenmiller, Peter Campbell, Piero Carninci, Menna Clatworthy, Hans Clevers, Bart Deplancke, Ian Dunham, James Eberwine, Roland Eils, Wolfgang Enard, Andrew Farmer, Lars Fugger, Berthold Göttgens, Nir Hacohen, Muzlifah Haniffa, Martin Hemberg, Seung Kim, Paul Klenerman, Arnold Kriegstein, Ed Lein, Sten Linnarsson, Emma Lundberg, Joakim Lundeberg, Partha Majumder, John C Marioni, Miriam Merad, Musa Mhlanga, Martijn Nawijn, Mihai Netea, Garry Nolan, Dana Pe’er, Anthony Phillipakis, Chris P Ponting, Stephen Quake, Wolf Reik, Orit Rozenblatt-Rosen, Joshua Sanes, Rahul Satija, Ton N Schumacher, Alex Shalek, Ehud Shapiro, Padmanee Sharma, Jay W Shin, Oliver Stegle, Michael Stratton, Michael J T Stubbington, Fabian J Theis, Matthias Uhlen, Alexander van Oudenaarden, Allon Wagner, Fiona Watt, Jonathan Weissman, Barbara Wold, Ramnik Xavier, Nir Yosef, and Human Cell Atlas Meeting Participants. The human cell atlas. Elife, 6, Dec 2017.

[26] H Robert Frost. Computation and application of tissue-specific gene set weights. Bioinformatics, Apr 2018.

[27] Matthew E Ritchie, Belinda Phipson, D. Wu, Yifang Hu, Charity W Law, Wei Shi, and Gordon K Smyth. limma powers differential expression analyses for rna-sequencing and microarray studies. Nucleic Acids Res, 43(7):e47, Apr 2015.

[28] Christopher R. Genovese, Kathryn Roeder, and Larry Wasserman. False discovery control with p-value weighting. Biometrika, 93(3):509–524, 2006.

[29] Di Wu, Elgene Lim, François Vaillant, Marie-Liesse Asselin-Labat, Jane E Visvader, and Gordon K Smyth. ROAST: rotation gene set tests for complex microarray experiments. Bioinformatics (Oxford, England), 26(17):2176–2182, September 2010. PMID: 20610611.

